# Multivariate EEG analyses support high-resolution tracking of feature-based attentional selection: Multivariate EEG tracks feature-based selection

**DOI:** 10.1101/082818

**Authors:** Johannes Jacobus Fahrenfort, Anna Grubert, Christian N. L. Olivers, Martin Eimer

## Abstract

The primary electrophysiological marker of feature-based selection is the N2pc, a lateralized posterior negativity emerging around 180-200 ms. As it relies on hemispheric differences, its ability to discriminate the locus of focal attention is severely limited. Here we demonstrate that multivariate analyses of raw EEG data provide a much more fine-grained spatial profile of feature-based target selection. When training a pattern classifier to determine target position from EEG, we were able to decode target positions on the vertical midline, which cannot be achieved using standard N2pc methodology. Next, we used a forward encoding model to construct a channel tuning function that describes the continuous relationship between target position and multivariate EEG in an eight-position display. This model can spatially discriminate individual target positions in these displays and is fully invertible, enabling us to construct hypothetical topographic activation maps for target positions that were never used. When tested against the real pattern of neural activity obtained from a different group of subjects, the constructed maps from the forward model turned out statistically indistinguishable, thus providing independent validation of our model. Our findings demonstrate the power of multivariate EEG analysis to track feature-based target selection with high spatial and temporal precision.

**Significance Statement:** Feature-based attentional selection enables observers to find objects in their visual field. The spatiotemporal profile of this process is difficult to assess with standard electrophysiological methods, which rely on activity differences between cerebral hemispheres. We demonstrate that multivariate analyses of EEG data can track target selection across the visual field with high temporal and spatial resolution. Using a forward model, we were able to capture the continuous relationship between target position and EEG measurements, allowing us to reconstruct the distribution of cortical activity for target locations that were never shown during the experiment. Our findings demonstrate the existence of a temporally and spatially precise EEG signal that can be used to study the neural basis of feature-based attentional selection.

## Introduction

Feature-based attention serves to select relevant input from competing information. This is most prominent during visual search, when observers look for target objects among distractors, and selective attention is guided on the basis of target-defining features^e.g. 1^. Neurophysiological markers of feature-based selection have been found in single-unit recording studies in nonhuman primates, where objects with target-matching features triggered increased neural responses at corresponding locations in retinotopic visual cortex^2,3^. In humans, EEG/MEG-based measures have provided the temporal resolution required to track feature-based selection in real time, but the spatial resolution of these measures is typically regarded as poor. Although the topography of early visual event-related potential (ERP) components varies with the location of a single visual stimulus in an otherwise empty visual field^4^, it is more challenging to discriminate the position of attended objects in multi-stimulus visual search displays using EEG/MEG markers.

One EEG/MEG signal in particular has become the gold standard of feature-based selection: The N2pc component of the event-related potential. The N2pc is an enhanced negativity elicited around 200 ms post-stimulus at posterior electrodes contralateral to candidate target objects^e.g. 5,6^. It is generated in extrastriate ventral visual cortex^7^, and is assumed to reflect the spatially selective enhancement of neural activity during feature-based target selection^see^ ^8,9^ ^for details^. Although N2pc components are widely used, their spatial resolution is severely limited. N2pc components are computed by subtracting ipsilateral from contralateral potentials, so the spatial information is limited to activation differences between cortical hemispheres. Although N2pcs are generally larger for the lower versus upper visual field^10^, they are not well suited to discriminate between attentional selection within the same hemifield or quadrant, and completely blind to the selection of targets on the vertical meridian (above and below fixation).

The present study tested the idea that the distribution of EEG activity across the scalp contains more precise spatial information than what is shown using the N2pc. To investigate this, we employed two types of multivariate analyses. *Backward decoding models* (BDMs) predict which stimulus condition (e.g., an attended target at a particular location) was present on the basis of EEG activity patterns. Above-chance classification accuracy then shows that the relevant information is represented in EEG signals, and the time course of classification performance indicates at which post-stimulus latency this information becomes available. BDM classifier outputs can also be used to construct maps showing the topography of the neural activity underlying successful classifications^11^. *Forward encoding models* (FEMs) reverse this direction of inference by describing the relationship between multivariate neural activity and an experimental variable that is hypothesized to be continuous in nature (here the attended location). The relationship between this variable and the multivariate signal can be described using a Channel Tuning Function (CTF)^12,13^. For example, multivariate analyses of alpha-band EEG activity have recently been employed to track the coding of spatial locations in working memory^14^ and during endogenously cued spatial attention shifts^15^. Here, we applied both BDMs and FEMs to raw EEG signals to extract and reconstruct feature-based target selection processes in space and in time.

In Experiment 1, colour-defined targets appeared on the horizontal or vertical midline in two successive displays (Figure 1). Both horizontal and vertical target positions were successfully classified using BDMs from around 200 ms after display onset, and with topographies consistent with the retinotopic architecture of early visual cortex. In Experiment 2, targets could appear at eight locations, but never on the vertical or horizontal midline (Figure 2). Forward modelling yielded Gaussian-shaped Channel Tuning Functions (CTFs), which we used to construct hypothetical channel responses and activation maps for targets placed on the horizontal and vertical midline. The resulting maps were then validated through the topographies observed in Experiment 1 (where these targets were actually presented). These results demonstrate the power of backward and forward modelling of multivariate EEG data in tracking attentional selection across both time and space.

**Figure 1.**
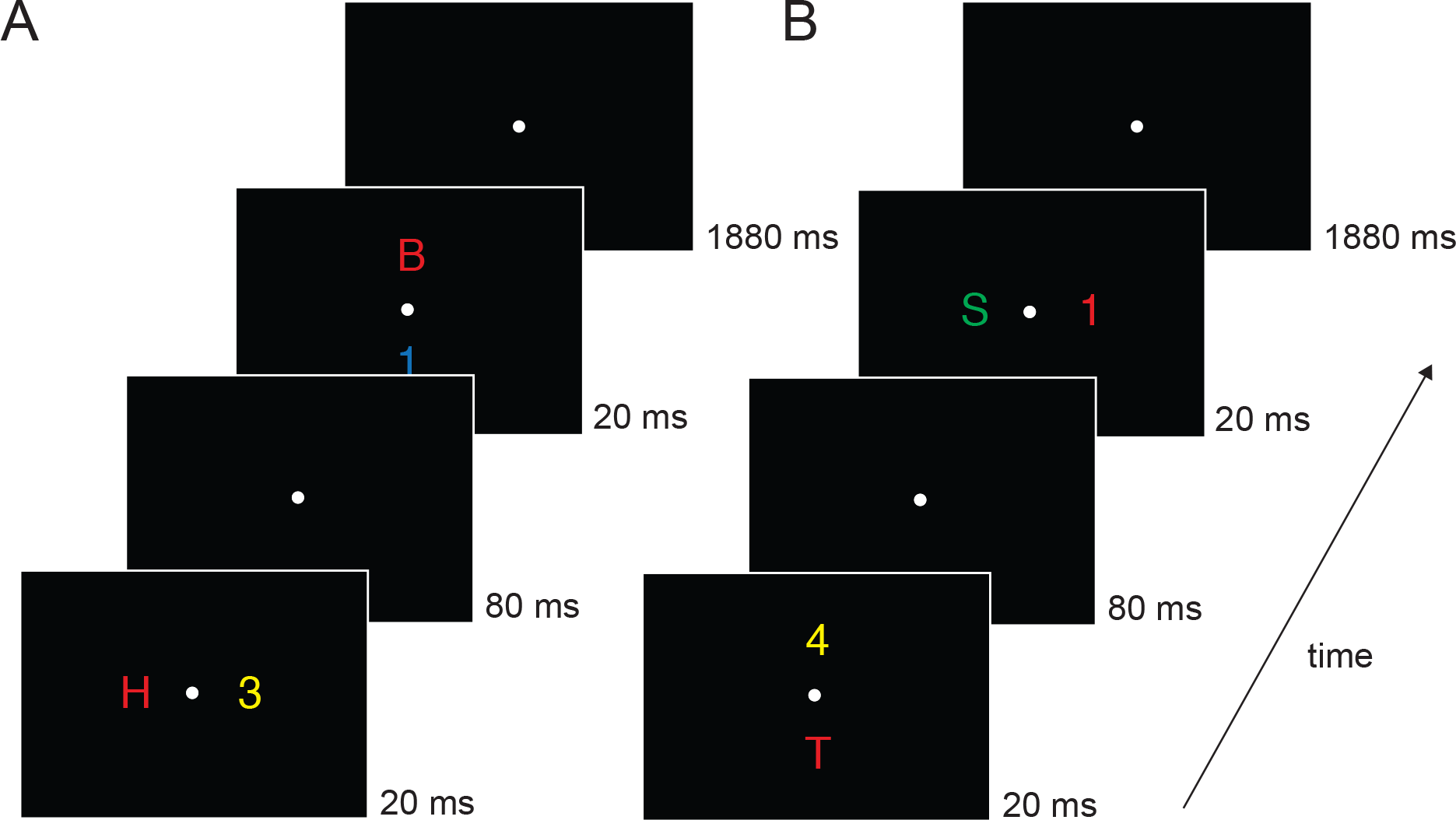
Example trial time lines of Experiment 1. There were two types of trials: the first display contained items on the horizontal axis and the second display items on the vertical axis ***(A)***, or the first display contained items on the vertical axis and the second display items on the horizontal axis ***(B)***. Subjects were required to determine whether a color-defined target was a digit or letter. The target color remained constant within a session (red in this example). Potential target colors were red, green, blue or yellow (counterbalanced across subjects). Within blocks, subjects either had to detect a target in the first display (D1 blocks) or they had to detect a target in the second display (D2 blocks). We only analyzed task relevant displays.

**Figure 2.**
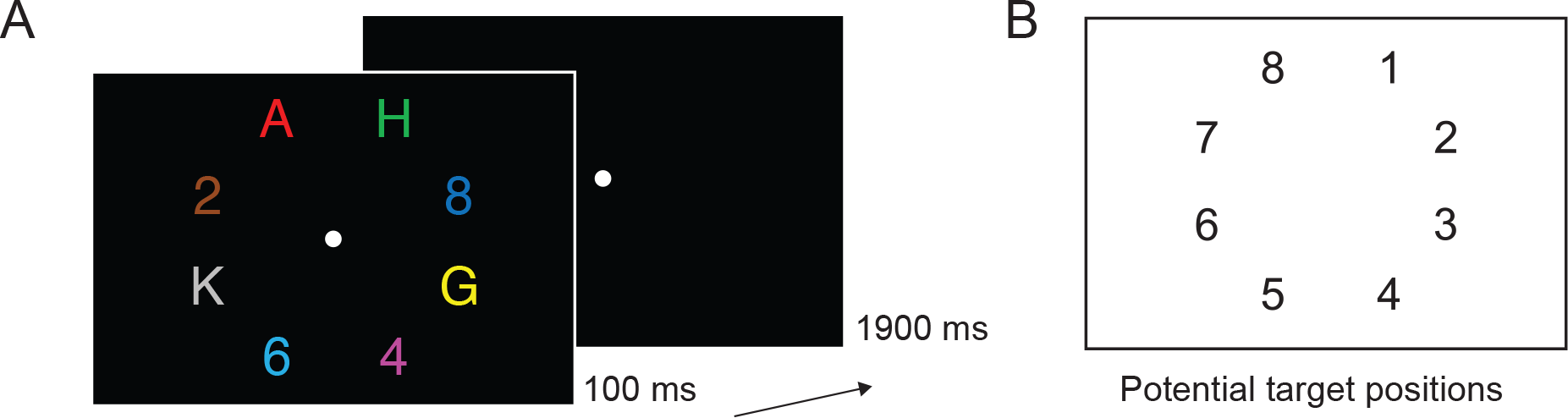
Task and conditions in Experiment 2. ***(A)*** Trial time line of Experiment 2. Subjects were asked to determine the identity of a coloured target (letter or digit), target colour could be red, green or blue (counterbalanced across subjects). ***(B)*** Positions used in Experiment 2, counted clockwise, one position contained the target, the other positions were occupied by distractors.

## Methods

### Participants

Fifteen subjects were paid to participate in Experiment 1. Three showed excessive eye movement activity (containing blinks and/or eye-movements on more than 60< of all trials) and were therefore excluded from further analysis. The remaining twelve participants were aged between 25 and 37 years (mean age 28.8 years); five of them were female; two were left-handed. Non-overlapping analyses of data from these subjects have been published elsewhere^16^. Fifteen paid subjects participated in Experiment 2 (aged 25 to 42 years; mean age 31.2 years). Ten were female; four were left-handed. All participants in both experiments had normal or corrected-to-normal vision, including normal colour vision, which was substantiated by means of the Ishihara colour vision test^17^. Both experiments were conducted in accordance with the Declaration of Helsinki and were approved by the Psychology Ethics Committee at Birkbeck (University of London). All participants gave informed written consent prior to testing.

### Equipment

All stimuli were presented on a 22-inch Samsung wide SyncMaster 2233 LCD monitor (1280x1024 pixel resolution, 100 Hz refresh rate) against a black background. Participants were seated in a dimly illuminated, soundproof and electrically shielded testing booth and viewed the screen at a distance of 100 cm. Manual responses were collected by means of two purpose-built response keys which were vertically aligned and centred in front of the participants. Stimulus presentation, timing, and response recording were controlled by a LG Pentium PC running under Windows XP, using the Cogent 2000 toolbox (www.vislab.ucl.ac.uk/Cogent) for MATLAB (Mathworks, Inc., USA).

### Stimuli and procedure Experiment 1

Each trial contained two successively presented stimulus displays (Figure 1), each of which was shown for 20 ms. The two consecutive stimulus arrays were separated by a 100 ms SOA (80 ms blank screen between displays). Each display contained two items on opposite sides, one in the target colour (e.g., red in Figure 1), and another one in a nontarget colour. Four possible stimulus colours were used: red (CIE colour coordinates .616/.338), green (.261/.558), blue (.183/.178), and yellow (.399/.476). All colours were equiluminant (∼11.8 cd/m2). Target colour was counterbalanced across participants so that always three participants searched for targets in one of the four possible colours. In each trial, two different nontarget colours were randomly chosen from the set of the three remaining nontarget colours. One of the two displays on each trial contained a target-nontarget colour pair on the horizontal midline (to the left and right of fixation), and the other display contained a target-nontarget colour pair on the vertical midline (above and below of fixation). Presentation sequence (vertical stimulus pair preceded by horizontal pair or vice versa) was randomised across trials. Stimuli were uppercase letters (B, H, S, or T) and digits (1, 2, 3, or 4), subtending 0.9° x 0.9° of visual angle, and presented at an eccentricity of 2.4° from central fixation (with respect to the centre of each stimulus). The identities of the four stimuli shown on each trial were randomly selected from the set of all letters and digits. A central grey (.324/.348) fixation point (0.2° × 0.2°) was present throughout each block, and participants were instructed to maintain fixation on this point.

In different blocks, participants were instructed to report the category of the target colour item (digit or letter) in the first display and ignore the target colour item in the second display (D1 blocks), or report the target in the second display and ignore the first display (D2 blocks). Subjects indicated the category of the target by pressing the corresponding response key (e.g. digit on top key, letter on bottom key). The hand-to-key mapping (left or right hand on top or bottom key) was counterbalanced across participants, but remained constant for each participant.

The experiment contained 12 blocks with six successive blocks in which either the target-colour item in the first display (D1 blocks) or in the second display (D2 blocks) was task relevant. Six participants completed the D1 blocks before the D2 blocks, and this order was reversed for the other six participants. Each block comprised 64 trials, resulting in a total of 768 experimental trials. On 32 trials of each block, the horizontally arranged stimulus set preceded the vertically arranged stimulus set and vice versa on the other 32 trials. Target location (left/right/top/bottom) was always unpredictable in both displays. One practice block was run prior to the first experimental block.

### Experiment 2

In each trial, participants viewed a search display containing eight items in eight different colours that were located on an imaginary circle with an eccentricity of 2.4° of visual angle from fixation (with respect to each object’s centre; Figure 2A). Four items appeared in the left and four in the right visual field (represented as positions 1-8, in a clockwise fashion, starting from the upper right position; Figure 2B). Search displays were presented for 100 ms, and were followed by a blank 1900 ms interstimulus interval. A central grey fixation point (0.2° × 0.2°) was continuously presented throughout each experimental block, and participants were instructed to fixate this point throughout. Display items (0.6° × 0.6° each) were capital letters (A, K, G, H, N, F, X, and Y) and digits (2, 3, 4, 5, 6, 7, 8, and 9). For each trial, eight of these items were randomly chosen (without replacement). The location and colour of each item was randomly assigned, with colours chosen from the set of red (CIE colour coordinates .612/.327), green (.286/.594), blue (.174/.148), yellow (.389/.519), cyan (.211/.309), magenta (.217/.109), brown (.505/.412), and grey (.288/.316). All colours were equiluminant (∼10.4 cd/m^2^). Although the size of all displays items was identical (0.6° × 0.6°), the number of pixels for each item was not exactly matched, which means that overall luminance values may have differed slightly between individual items. However, because target and non-target items and the position of target items were selected randomly on each trial, no systematic luminance-related asymmetries associated with target location were present across trials. Participants’ task was to find the item in a pre-specified target colour and report its category (letter or digit) by pressing the corresponding upper or lower response key. Category-to-key and hand-to-key mappings were counterbalanced across participants. Red, green or blue each served as target colour for 5 participants, and this target colour remained constant for each participant throughout the experiment. Target position (1 to 8) varied randomly and unpredictably across trials.

The experiment comprised 12 blocks of 64 trials, resulting in a total of 768 experimental trials. Each block contained 4 trials for each combination of target position (1 to 8) and target identity (digit, letter). A practice block was run prior to the first experimental block.

### EEG recording and preprocessing

In both experiments, continuous EEG was DC-recorded using the BrainVision recorder software and a BrainAmp DC amplifier (Brain Products, Munich, Germany). Sampling rate was 500 Hz (sampling interval 2000 µS), with a resolution of 0.1 µV. The upper cut-off frequency during DC recording was 250 Hz, and EEG was digitally low-pass filtered at 40 Hz. In Experiment 1, EEG was recorded from 23 scalp sites at Fpz, F7, F3, Fz, F4, F8, FC5, FC6, T7, C3, Cz, C4, T8, CP5, CP6, P7, P3, Pz, P4, P8, PO7, PO8, Oz. In Experiment 2, four additional posterior recording sites were included (P9/P10 and PO9/PO10). The continuous EEG was sampled at 500 Hz and digitally low-pass filtered at 40 Hz during acquisition. All electrodes were online referenced to the left earlobe. Impedances were kept below 5kΩ.

EEG was re-referenced offline to the average of both earlobes and highpass-filtered using a Hamming windowed sinc FIR filter at 0.1 Hz (cutoff frequency .05 Hz) with a length of 33 seconds (pop_eegfiltnew.m from the EEGLAB toolbox), to remove slow drifts from the signal. Trials containing incorrect responses or responses occurring after 1500 ms were removed. On average 5.58< (SD 4.26) of all trials were removed in Experiment 1, and 2.66< (SD 1.54) of trials were removed in Experiment 2. In Experiment 1, task-irrelevant displays elicited reliable N2pc components only in D2 blocks where they preceded task-relevant displays but not in D1 blocks where they appeared after the task-relevant display was shown (as reported in detail elsewhere15). For this reason, epochs were extracted only for the task-relevant display (the first display in D1 blocks and the second display in D2 blocks), from -200 ms to 400 ms relative to the onset of the target display, corrected relative to a -200 to -100 ms pre-stimulus baseline. This particular baseline window was chosen for Experiment 1 because task-relevant displays in D2 blocks were preceded by another task-irrelevant display (with an SOA of 100 ms), in order to ensure that the baseline periods for epochs from D2 blocks would not be contaminated with EEG activity triggered by the presentation of the first display. For Experiment 2, data was segmented into -100 to 1000 ms windows, using the -100 ms to 0 ms period for baseline correction. All EEG was downsampled to 250 Hz and no further pre-processing was applied.

Stimulus presentation durations were too short to allow for meaningful saccades to the target in time, and so our main analyses therefore did not control for eye-movements. However, to investigate the potential influence of eye blinks and eye movements, we also performed our analyses on cleaned data. We removed eye-blink components using ICA, and rejected eye-movements after blink removal using a step-algorithm on the Horizontal Electro-Oculogram (HEOG). HEOG was obtained by taking the difference between the electrode on the left and the right outer canthi of the eyes. The step algorithm used a 100 ms sliding window with steps of 50 ms in the period 0-500 ms after stimulus presentation, identifying amplitude fluctuations of more than 25 μV as eye-movements. This procedure was performed on the data from both experiments, confirming that very few eye-movements occurred in either experiment (Experiment 1: 0.46<, SD 0.71<, Experiment 2: 0.99<, SD 1.35<). The data with blink-removal and eye-movement rejection yielded statistically indistinguishable classification accuracies and CTFs when compared to data with eye blinks and eye-movements included.

### N2pc analyses

For each subject in Experiment 1, event-related potentials (ERPs) to task-relevant target items (first display targets in D1 blocks, and second display targets in D2 blocks) were computed separately for targets to the right and to the left of fixation, collapsing over target identity (digit or letter) and display position (first or second display). ERP averages were balanced to contain an equal number of digit and letter targets as well as an equal number of targets from the first and second display. N2pc components were computed for ERPs at lateral posterior electrode sites PO7 and PO8 by subtracting ERPs recorded at the electrode ipsilateral to the target location from contralateral ERPs. Statistical tests were performed on the resulting contralateral-ipsilateral N2pc difference waveforms, using two-sided t-tests against zero for each time sample, and were corrected for multiple comparisons using cluster-based permutation testing on contiguously significant samples using 1000 iterations^18^. For Experiment 1, no N2pc components could be computed for target objects at the top versus bottom positions, because the N2pc is defined as the difference between contralateral and ipsilateral posterior ERPs in response to attended objects in the left and right visual field. This standard logic of subtracting ipsilateral from contralateral ERP waveforms cannot be applied to search displays that contain a non-lateralized target objects on the vertical meridian. For Experiment 2, N2pc components were measured for targets in the left versus right visual field, averaged across all four possible target positions on either side. In addition, separate N2pc components were computed for left versus right target objects at each of the four vertical elevations.

### Backward Decoding Model (BDM)

For each participant in both experiments, we applied a backward decoding classification algorithm, using a 10-fold cross validation scheme. First, we removed information about the order in which trials were acquired during the experiment by randomizing the order in which trials were stored on disk. Next, we split up the dataset into 10 equally sized subsets. Subsequently, a linear discriminant classifier was trained to discriminate between stimulus classes using 90% of the data, and was tested on the remaining 10% of the data, thereby ensuring independence of training and testing sets. This procedure was repeated 10 times until all data had been tested once. The EEG amplitudes at individual electrodes were used as features for classification, resulting in 23 features per stimulus class in Experiment 1 and 27 features in Experiment 2. Classification accuracy for each subject was computed as the average number of correct class assignments, first separately for each condition, then averaged across conditions, and finally averaged across the 10 folds. This cross-validation procedure was executed for every time sample in a trial, yielding the evolution of classification accuracy over time.

For Experiment 1, separate classifications were performed for trials with horizontal targets (computing accuracy for discriminating between left versus right targets) and for trials with vertical targets (computing discrimination accuracy for top versus bottom targets). The same trial sets as in the N2pc analysis were used. For Experiment 2, we performed two BDM analyses. In the first analysis, we entered the eight target positions into the classifier as eight separate stimulus classes. This 8-way classification was performed to determine whether the classifier could discriminate between target positions across the visual field. In the second analysis, we conducted four separate sets of classifications for trials with target objects within each of the four quadrants (positions 1 versus 2; 3 versus 4; 5 versus 6; 7 versus 8), to test whether target positions could be also be reliably discriminated within quadrants.

Statistical tests were performed across subjects using a two-sided t-test against chance for each time sample, correcting for multiple comparisons using group-wise cluster-based permutation testing on contiguously significant (p<.05) time samples. This procedure takes the sum of the t-values for all contiguously significant time points, and computes the number of times this sum is exceeded when computing the maximum cluster-based sum under random permutation (for details see ^18^). Although testing against chance is common, one caveat when using t-statistics on k-fold classification accuracy is that this does not allow population-level inference, in fact producing fixed-effects rather than random-effects results (see ^19^ for details). The implication is that one cannot formally draw population-level inferences based on such analyses, restricting conclusions to the sample that was tested. For studies that require population-level inference, it would be preferred to either use a completely separate training set (performing the training on different subjects or obtaining training data from a different task), or to replace the t-test with a statistic that explicitly evaluates information prevalence across the sampled subjects (again, see ^19^ for details).

To achieve temporal smoothing / remove high-frequency noise for presentation purposes only, the plotted BDM classification time series were fitted with a spline that was centered on maximum classification accuracy. This was achieved by downsampling the signal to 32 Hz, taking peak classification accuracy as the point of origin from which downsampling was applied (effectively centering the resampled data on peak classification accuracy). This was done to make sure that the height of the peak was not diluted by the smoothing procedure. Next, a spline was fitted through the data points that remained after downsampling. However, all statistics and visualizations of significant time windows were based on the original unfiltered data.

In addition to assessing classification accuracy across time, we also computed topographical maps based on classifier weights for individual features (electrodes). Because weights from backward decoding models cannot be reliably interpreted as neural activity, we obtained these maps by using a method recently described by Haufe and colleagues^11^, in which the classifier weights are multiplied with the data covariance matrix. Because the weights obtained from linear discriminant analysis contain the difference between the two compared sets normalized by the covariance matrix, this operation creates activation patterns that return the mass-univariate difference between the compared conditions, but which unlike classifier weights, are interpretable as neural sources. To be able to compare maps from Experiments 1 and 2, we spatially normalized each subjects’ activation pattern across electrodes by subtracting the mean across electrodes and dividing by the standard deviation across electrodes. As a result, topographic maps for which between-experiment comparisons are made represent Z-scores with average 0 and unit standard deviation, and show the normalized spatial distribution of EEG activity that underlies successful discrimination between stimulus classes.

### Forward Encoding Model (FEM)

While the BDM analyses used a classifier on multivariate EEG data to determine class membership for a fixed set of target positions, the FEM model reverses this direction of inference by capturing the continuous relationship between target positions and the multivariate EEG data. We employed a procedure previously described by Brouwer and Heeger^12^ using the same 10-fold cross validation scheme as in the BDM. In this procedure, the training set is used to model the response in each of eight hypothetical position “channels” (corresponding to the eight target positions on the screen). The nomenclature “channels” here should not be confused with MEG or EEG sensors, EEG sensors are referred to as electrodes in the current manuscript. The modelled response is called the “basis set”, which in our primary analysis contained a 1 for the corresponding target position and a 0 for all other positions (such a binary on-off response is sometimes referred to as a delta function). We used eight basis sets (one for each target position), each shifted by one position compared to its neighbour, to construct a regression matrix C1. C1 has the form k × n1, in which k is the number of position channels (1 to 8) and n1 is the number of trials in the training set. Next we estimated the response amplitude to each of the eight hypothetical position channels by performing an ordinary least squares regression of the C1 matrix onto the B1 matrix from the EEG training set. B1 contained EEG data of the form m × n1, in which m is number of electrodes and n1 is the number of trials in the training set. This regression yields a weight matrix W in which each electrode obtains a regressor weight for each target position. The weight matrix W has the form m × k, in which m is the number of electrodes and k is the number of position channels.

Next, the model is inverted by performing ordinary least squares regression of these weights onto the B2 matrix from the EEG testing set to produce the estimated channel responses for each trial. B2 has the form m × n1, in which m is number of electrodes and n2 is the number of trials in the testing set. The resulting estimated channel responses are contained in matrix C2, having the form k × n2, in which k are the observed channel responses and n2 are the trials in the testing set. This procedure is repeated for all folds in the train-test procedure, until all data has been tested once. Next, the channel responses are averaged across trials in the testing set, separately for each of the eight trial types that correspond to each of the eight target positions on the screen. The channel outputs from this testing phase in combination with the associated weights contain the validated and invertible one-to-one relationship between a particular attended location in the search display and the multivariate EEG response. Finally, the estimated channel responses in C2 are shifted to a common center (channel position 4, in this case), so that the channel responses for each of the eight target positions are aligned (see Fig. 8B for the individual target-position CTFs, repeating the most distant position to achieve symmetry for display purposes) and then again averaged across trial type to generate the empirical canonical CTF (see Fig. 8A for the averaged CTF). This entire procedure is repeated for every time point, yielding a CTF over time (see Fig. 8C). A mathematical (less verbal) description of this training-testing FEM estimation procedure has been provided elsewhere, both for fMRI, MEG and EEG data^e.g. 12-15,20,21^.

For the CTF time series plots, we applied spline interpolation over both channels and time to achieve smoothing for presentation purposes only. All statistical tests and visualizations thereof were based on the original data. In addition, we characterized the topography of estimated neural activity associated with allocating attention to specific target locations by plotting the corresponding regression weight matrix, separately for each of the 8 conditions. Weight maps from forward models are meaningful because regression weights in forward models are directly interpretable in terms of neural sources^11^.

Most studies applying a forward modelling approach use a basis set that assumes a particular shape of the CTF by adopting a hypothetical model for the relationship between variations in stimulus space and the multivariate response. However, because the width and shape of this basis set cannot be known beforehand, such assumptions can have an impact on the weights that are obtained from the training data, and may even overestimate the Gaussian nature of the underlying CTFs. The alternative approach that we employed here is to not make any assumption about the shape of the CTF, and simply set the channel response to 1 for each target position, while setting all the other channel response to 0 (delta function)^e.g. also see 13^. This makes it possible to characterize the ‘real’ shape of the CTF using the testing data without making any a priori assumptions. Observing a continuous (i.e., Gaussian-shaped) CTF profile under these conditions guarantees that this shape is a dominant feature of the underlying dataset. Importantly, such a continuous response profile would imply that a CTF accurately describes the relationship between neural activity and a continuous stimulus property (target position in the present study). In this case, it is possible to predict the multivariate response pattern for target positions that were not actually presented in a particular experiment. Based on this logic, we tested whether the CTFs observed in Experiment 2 where targets were never presented on the vertical or horizontal midline could be used to reconstruct the pattern of neural activity that was observed in Experiment 1 for targets that appeared at these positions (see Results section for details).

### Analysis software

All data were analyzed in Matlab (the Mathworks, Inc., USA). Preprocessing was done using the EEGLAB toolbox^22^. All BDM and FEM analyses and visualizations were performed using a custom written toolbox: the Amsterdam Decoding and Modelling (ADAM) toolbox that uses EEGLAB or FieldTrip as input format. This custom written toolbox contains a general-purpose set of functions for BDM and FEM analyses, statistics and visualizations on group EEG data. The toolbox will also be released to the public domain, but until then the scripts are available upon request from the corresponding author.

## Results

### Experiment 1: Behavioral results

Trials with slow (> 1500 ms) reaction times (RTs) were excluded from analysis (fewer than 1% of all trials). Mean correct RTs as well as error rates were very similar for trials in which participants responded to a horizontally (604 ms; 5.0 %) or vertically aligned target (613 ms; 5.9 %), both *t*(11) < 1.9, *p* > .097.

### Experiment 1: N2pc and BDM results

As expected, a reliable N2pc component was elicited for task-relevant stimulus displays where the colour-defined target object was presented on the left or right side (Figure 3). This N2pc first reached significance at 164 ms after stimulus onset, and remained significant for about 100 ms (two-tailed cluster p-value = .003). The later contralateral negativity that followed the N2pc at around 300 ms after stimulus onset (see Figure 3, two-tailed cluster p-value = .003) has also been observed in previous ERP studies of visual attention^23,24^. This sustained posterior contralateral negativity (SPCN) component is assumed to reflect the encoding of target stimuli in visual working memory that follows their initial selection^see 9,16^ ^for details^.

**Figure 3.**
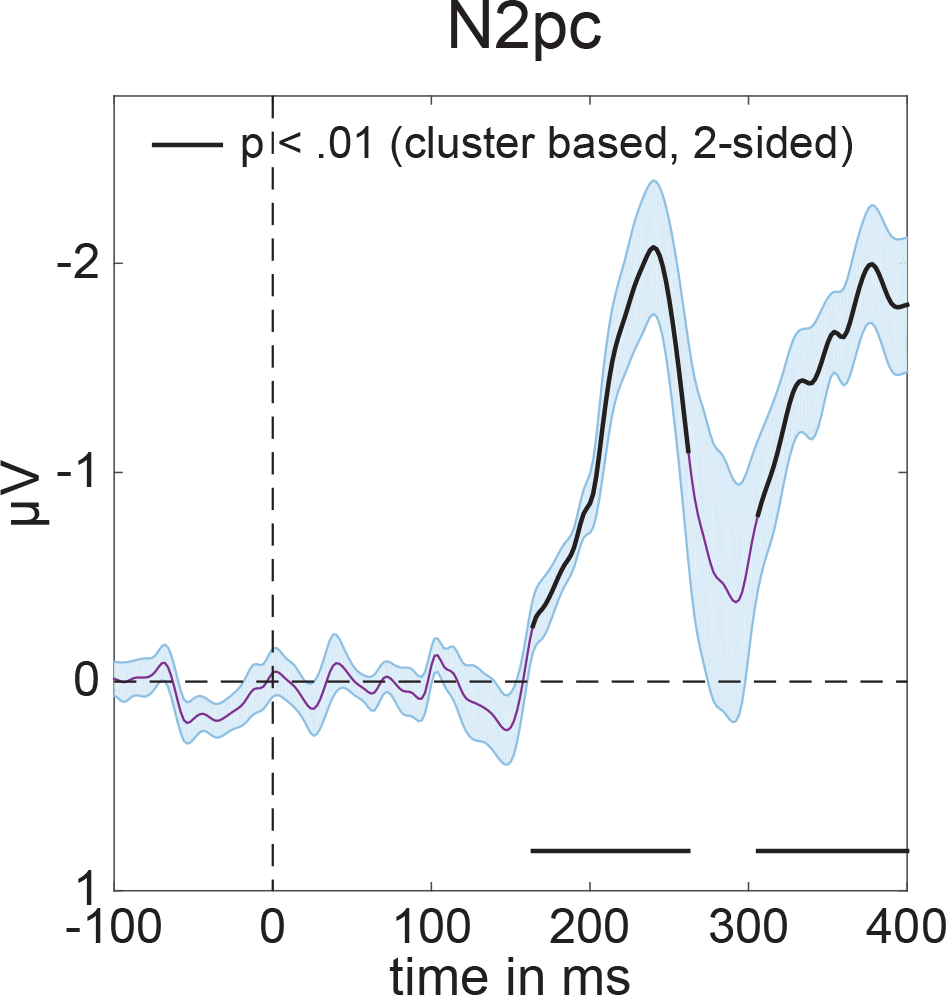
Average N2pc for left and right targets in Experiment 1. Computed using PO7 and PO8 (see methods). Black lines have p<.01 using cluster based permutation testing, see main text for exact p-values. Shaded areas show +/- s.e.m. Peak at 240 ms.

Because the N2pc is computed by subtracting ipsilateral from contralateral ERPs, it can only be measured for horizontally lateralized targets in the left versus right hemifield, but not for target objects that are presented on the vertical meridian (above versus below fixation). The central aim of Experiment 1 was to find out whether neural signals associated with the attentional selection of these target objects can be uncovered using multivariate pattern analysis. We trained a linear discriminant classifier to either discriminate whether targets appeared on the left or on the right (for targets on the horizontal axis), or at the top or the bottom (for targets on the vertical axis), based on the EEG responses across the scalp (see Methods for details). The classification accuracy results in Figure 4 show that this backward decoding model was not only able to discriminate left versus right targets (Figure 4A, two-tailed cluster p-value < 10^-3^ after 1000 iterations), but also successfully classified targets at the top versus bottom position (Figure 4B, two-tailed cluster p-value < 10^-3^ after 1000 iterations). The time course of classification accuracy was very similar for these two types of discriminations, and closely matched the time course of the N2pc plus SPCN components to lateral targets shown in Figure 3. From approximately 200 ms after the onset of a stimulus display, the ability of the classifier to discriminate between left/right and top/bottom targets increased rapidly, and classification performance remained above chance for the rest of the 400 ms time window analysed here. The successful classification of attended targets at top versus bottom positions during this time period demonstrates that non-lateralized information about the focus of attention at these positions was clearly present in the EEG signals, where this information remains hidden to the classical N2pc methodology.

**Figure 4.**
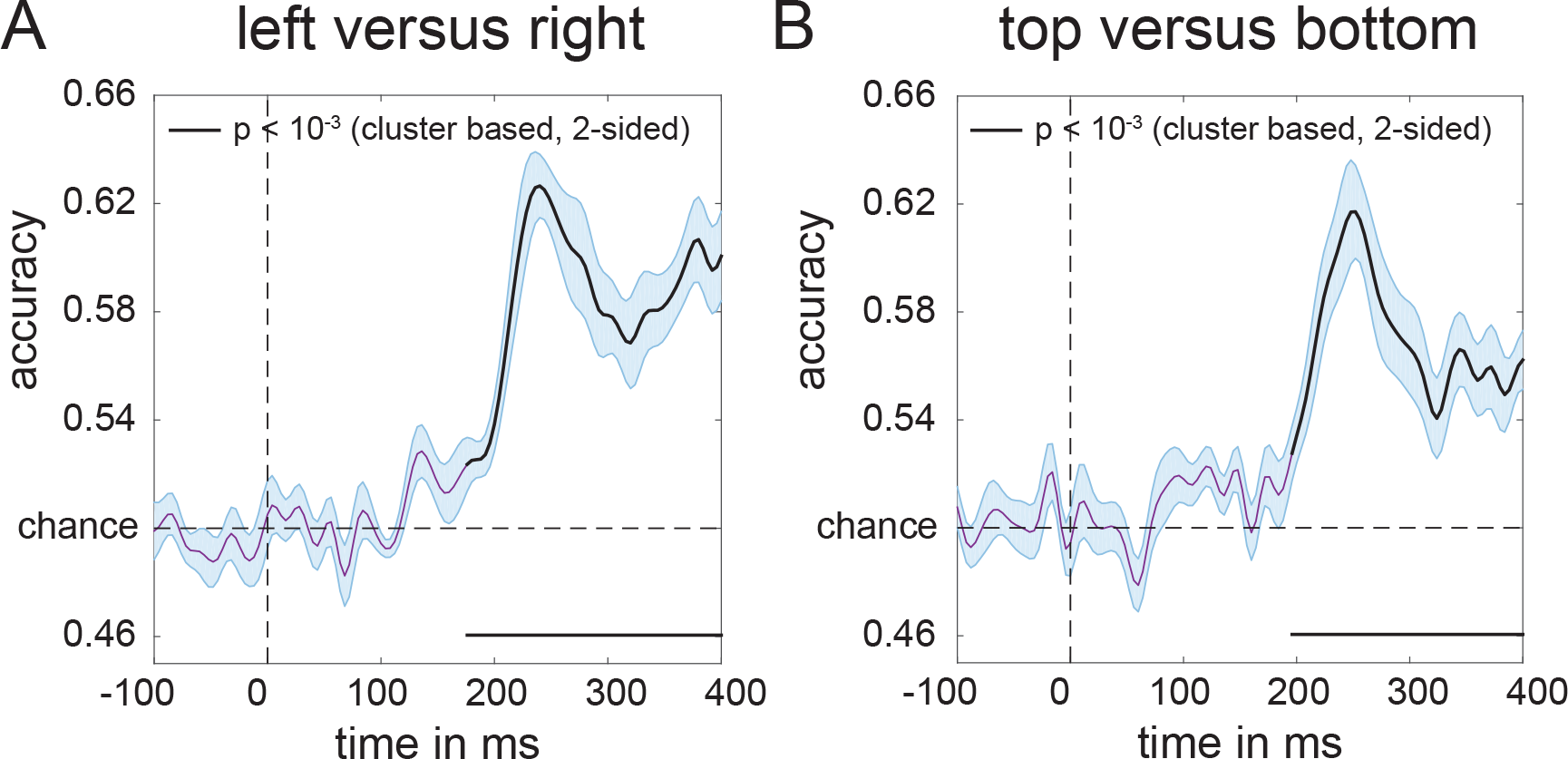
Classification accuracy of target position over time in Experiment 1. ***(A)*** Classification accuracy for left versus right targets. ***(B)*** Classification accuracy for top versus bottom targets. Black lines have p<10^-3^ using cluster based permutation testing. Shaded areas are +/− s.e.m. Peaks respectively at 244 ms (left-right) and 248 ms (top-bottom). Note the similar temporal evolution between N2pc (Figure 3) and classification accuracy here.

Next, we characterized the sources underlying these classifications by multiplying the weight matrix obtained from the classification analysis with the covariance matrix11 (see methods for details). The resulting topographical maps for left versus right and top versus bottom discriminations shown in Figure 5 represent the distribution of the neural signals contributing to these two types of classifications. They show a clear left-right gradient for left-right target discrimination (Figure 5A, two-tailed cluster p-values after 1000 iterations are p = .002 and p < 10^-3^ for the positive and negative cluster respectively), and a top-bottom gradient for top-bottom target discrimination (Figure 5B, p = .014 and p = .003 for the positive and negative cluster respectively), in line with the known representation of these locations in retinotopic visual cortex.

**Figure 5.**
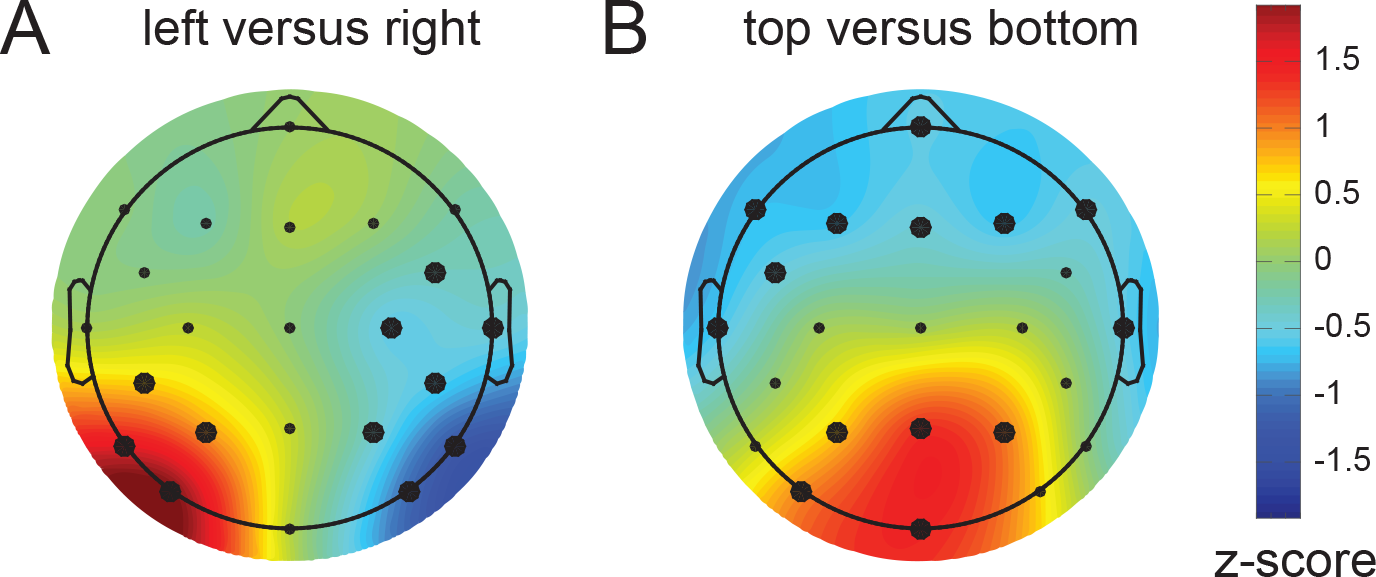
Activation patterns associated with peak decoding accuracy (240-250 ms) in Experiment 1. Derived from the product of the weight vectors and the covariance matrix (see Haufe et al, 2014) normalized across space (see Methods). ***(A)*** The pattern associated with left versus right decoding. Note the clearly lateralized distribution. This lateralized pattern shows the distribution of neural activity underlying successful discrimination between targets appearing on the left and the right of fixation, and is equivalent to the mass-univariate difference between left and right targets. ***(B)*** The pattern associated with top versus bottom decoding, now showing a posterior-anterior distribution. Thick electrode dots belong to clusters having p<0.05 under cluster based permutation testing, see main text for exact p-values.

### Experiment 2: Behavioral results

All responses occurred within 1500 ms after target display onset. There was a main effect of target side on reaction times, *F*(1,14) = 26.3, *p* < .001, 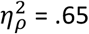, with faster responses for targets on the right (567 ms) as compared to the left display side (586 ms). Error rates were generally low (2.7%), and did not differ between left versus right targets.

### Experiment 2: N2pc and BDM results

All colour-defined target objects in Experiment 2 were presented in the left or right visual field, and produced reliable N2pc components within the same time window as the N2pc observed in Experiment 1. This is shown in Figure 6A, averaged across all target positions (two-tailed cluster p-value < 10^-3^ after 1000 iterations). Results of the multivariate BDM analysis of the EEG data based on all 8 possible target locations are shown in Figure 6B. This omnibus 8-way classification yielded results that were very similar to the time course of the N2pc component, with above-chance classification performance from about 190 ms post-stimulus (two-tailed cluster p-value < 10^-3^ after 1000 iterations). However, whereas the average N2pc only reflects neural response asymmetries associated with targets being presented in the left versus right visual field, the 8-way BDM analysis also takes variations of target position along the vertical axis into account. Some of these variations are also reflected by N2pc amplitudes, when N2pc components are computed separately for left versus right targets at different vertical elevations (positions 8 versus 1, 7 versus 2, 6 versus 3, and 5 versus 4, as shown in Figure 6C). N2pc amplitudes were generally larger for targets in the lower versus upper visual field, in line with previous observations^9^, and also for targets at more peripheral versus medial positions. However, and in contrast to multivariate BDM analyses, the N2pc components shown in Figure 6C are still computed on the basis of contralateral/ipsilateral differences in response to left versus right targets, and therefore cannot discriminate neural responses to target stimuli on the same side. Similarly, the N2pc is not able to fully discriminate between targets at different elevations, as for example the Upper and Bottom field presentations (see Figure 6C). To further underline this point, we assessed whether target locations can be successfully discriminated with BDM even when they are in the same visual quadrant. Figure 7 shows four additional BDM classifications, one for each quadrant. The comparisons were conducted for targets in the top right (positions 1 versus 2), bottom right (positions 3 versus 4), bottom left (positions 5 versus 6), and top left quadrants (positions 7 versus 8). As shown in Figure 7, all of these within-quadrant classifications were performed with above-chance accuracy for EEG data recorded between 200 – 300 ms post-stimulus (two-tailed cluster p-values for the first significant cluster after trial onset were p = .022 for top right, p = .001 for top left, and p < 10^-3^ for bottom right and bottom left). Accuracies were slightly lower than for the top-bottom and left-right discriminations in Experiment 1 (cf. Figure 4), plausibly due to the more fine-grained nature of the discriminations and a lower signal-to-noise ratio due to the lower number of trials for these comparisons when compared to Experiment 1.

**Figure 6.**
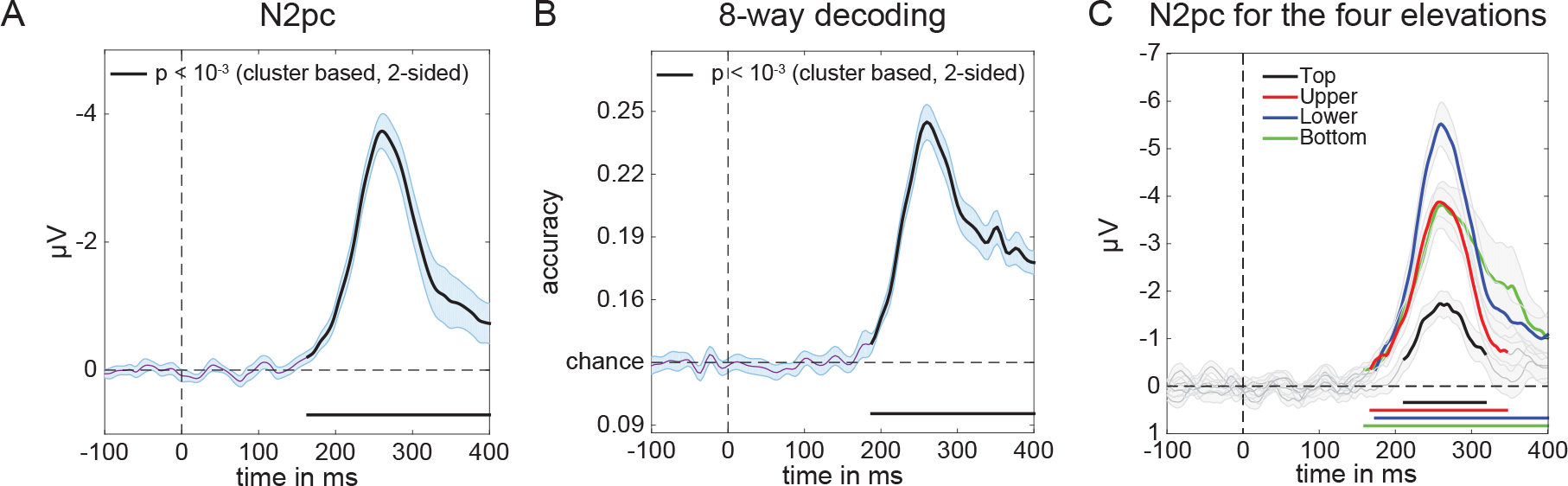
Decoding accuracy in Experiment 2. ***(A)*** 8-way classification accuracy of target position (chance = .125). The black line reflects significance at p<10^-3^ (cluster based permutation test), shaded area is +/− s.e.m. Peak at 264 ms. Note again the similarity in temporal evolution with the N2pc (figure 3) and decoding accuracies of Experiment 1 (Figure 4), although plausibly peaking slightly later because of the increase in the number of items on the screen. ***(B)*** Classsification accuracy per quadrant (chance = .5). ***(C)*** Separate N2pc components were computed for left versus

**Figure 7.**
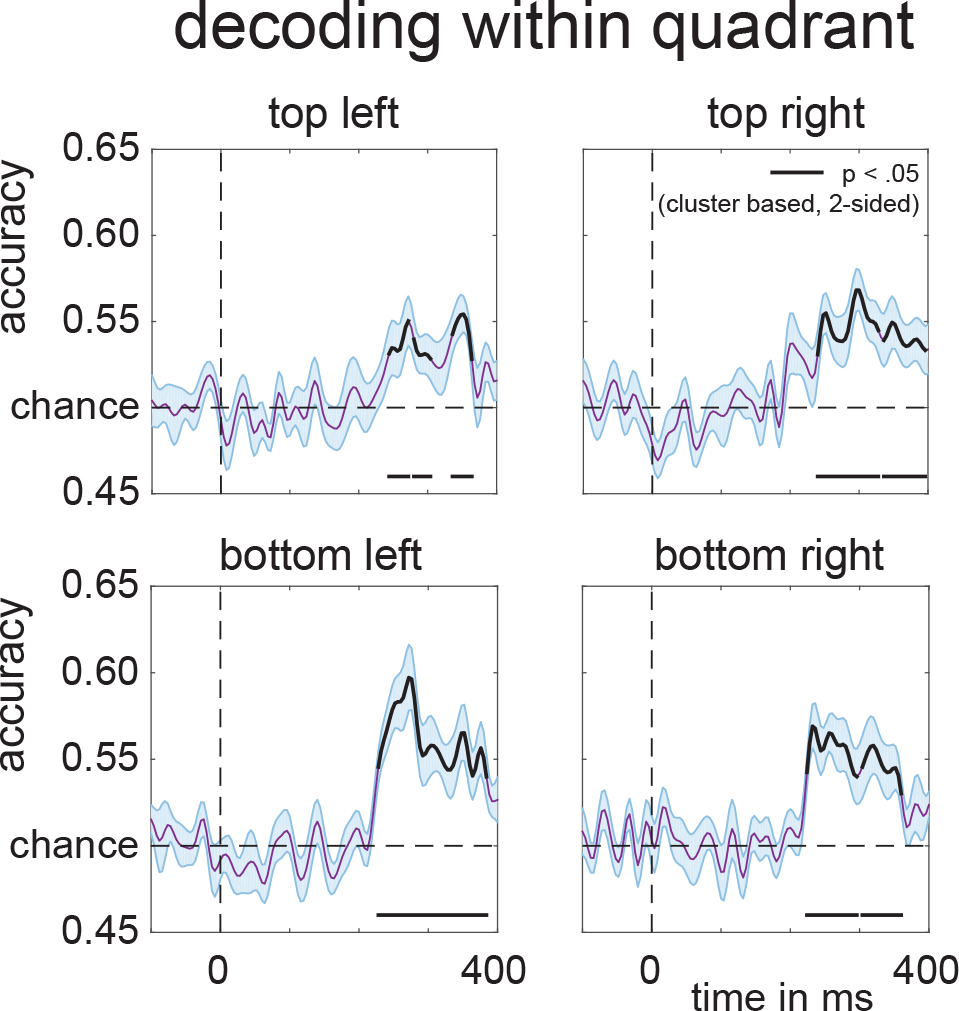
Per quadrant decoding accuracy of target position in Experiment 2. Black lines are corrected for multiple comparisons at p<.05 using cluster based permutation testing, two-tailed, see main text for exact cluster p-values. Shaded areas are +/− s.e.m.

### Experiment 2: FEM results

The ability to make more fine-grained discriminations between target positions within hemispheres and quadrants inspired us to build a continuous model of the relationship between target position and the multivariate EEG signal. For this we employed procedures similar to those used by Brouwer and Heeger^12^ and Garcia et al.^13^ (see Methods section for details). A clear channel tuning function (CTF) was obtained during the 260-270 ms post-stimulus time window (corresponding to the post-stimulus latency when BDM accuracy was maximal; see Figure 6B), describing a graded continuous relationship between attended target positions and multivariate EEG responses (Figure 8A). To determine whether this relationship was driven more strongly by some positions than by others, we also plotted the channel responses separately for each of the 8 stimulus conditions (Figure 8B). Although slightly more noisy, these results confirm that a Gaussian-like response profile was present for all 8 target positions, such that neighboring positions result in partially overlapping activation patterns. This confirms that the overall CTF was driven jointly by all target positions and not just by a subset of positions.

**Figure 8.**
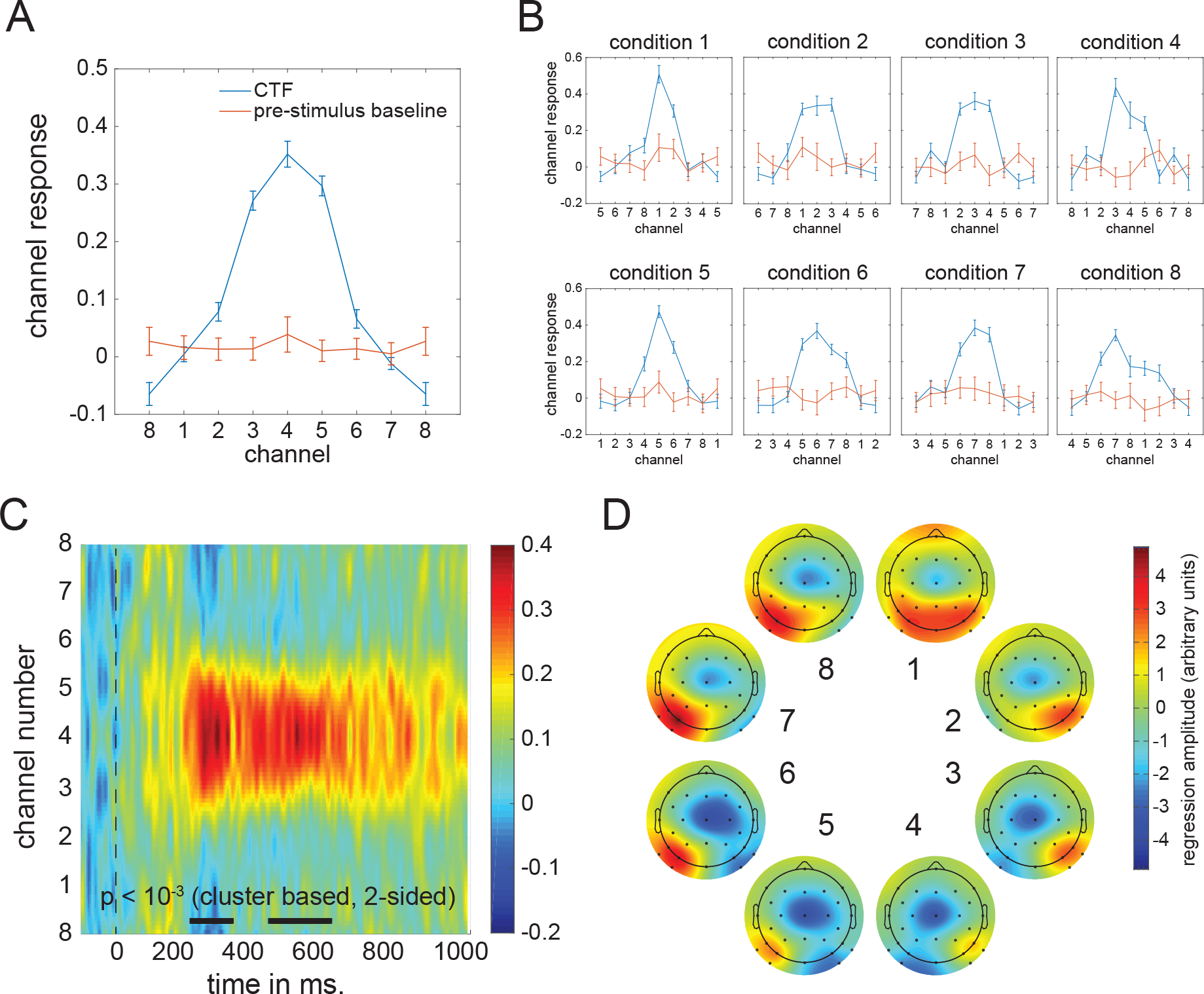
Channel Tuning Functions (CTF). ***(A)*** CTF for the 260-270 ms window and CTF during baseline (-100 to 0 ms) obtained by shifting the individual condition CTFs to align to the same channel. ***(B)*** CTFs for individual conditions show that the CTF is not driven by particular target positions. ***(C)*** CTF development over time in which color reflects channel responses. The black line near the time axis shows the time windows where the center position channel has significantly stronger channel responses than the outer boundary channel (p<10^-3^, cluster based permutation test). ***(D)*** Topographic weight plots for each condition in the 260-270 ms time window. Weights from forward models are directly interpretable in terms of neural sources (Haufe et al., 2014). These plots therefore show how neural activity changes as a function of variability in target position.

To determine how the CTF signal reflecting target position evolves in real time, we plotted the CTF as a function of time during a 1000 ms post-stimulus time window as a 2D plot in which colour reflects the strength of the channel response (Figure 8C). Statistical significance of the CTF was established by t-testing the center channel response to the channel response occurring in the periphery for each sample point, correcting for multiple comparisons using cluster-based permutation testing. The thick black line near the time axis represents statistically significant periods (both p < 10^-3^ after 1000 iterations). These plots confirm that the strongest CTF for target locations in the 8-item search displays was present in the 200-300 ms time range, in line with the decoding accuracy results shown in Figure 6.

To further characterize the distribution of neural activity associated with particular target positions within the search displays, Figure 8D shows topographic plots of the electrode weights resulting from the FEM estimation procedure, which are directly interpretable in terms of neural sources^11^. The resulting plots show gradual and systematic topographical shifts while spatial attention moves from upper to lower target positions on the same side, and a clear polarity swap when attention moves from the right to the left side in (positions 1 to 4 versus positions 5 to 8).

### Experiment 2: Using FEM to construct activation patterns for new target positions

The other main goal of Experiment 2 was to perform an ultimate validity check of the FEM approach by testing whether it is possible to predict the distribution of neural responses to target positions that were not actually presented. In Experiment 2, all stimulus positions were lateralized and targets never appeared on the vertical or horizontal midline. Forward models that produce a graded CTF (as shown in Figure 7) can essentially be regarded as continuous models. The CTF obtained for Experiment 2 should therefore be able to predict hypothetical activation patterns in response to target locations that were not included in this experiment, such as the horizontal and vertical positions that were only employed in Experiment 1. To construct virtual activation patterns for vertical and horizontal target positions for Experiment 2, we used a line symmetrical version of the group CTF from Figure 8A as the basis set to capture the relationship between multivariate responses and target positions. By averaging the responses across two neighboring positions (i.e., by interpolating the channel responses between position 8 and 1 to obtain the hypothetical response to targets at the top location), new channel responses were reconstructed that correspond to the intermediate positions between 8 and 1 (top), 4 and 5 (bottom), 2 and 3 (right), and 6 and 7 (left; see Figures 9A and 9B).

**Figure 9.**
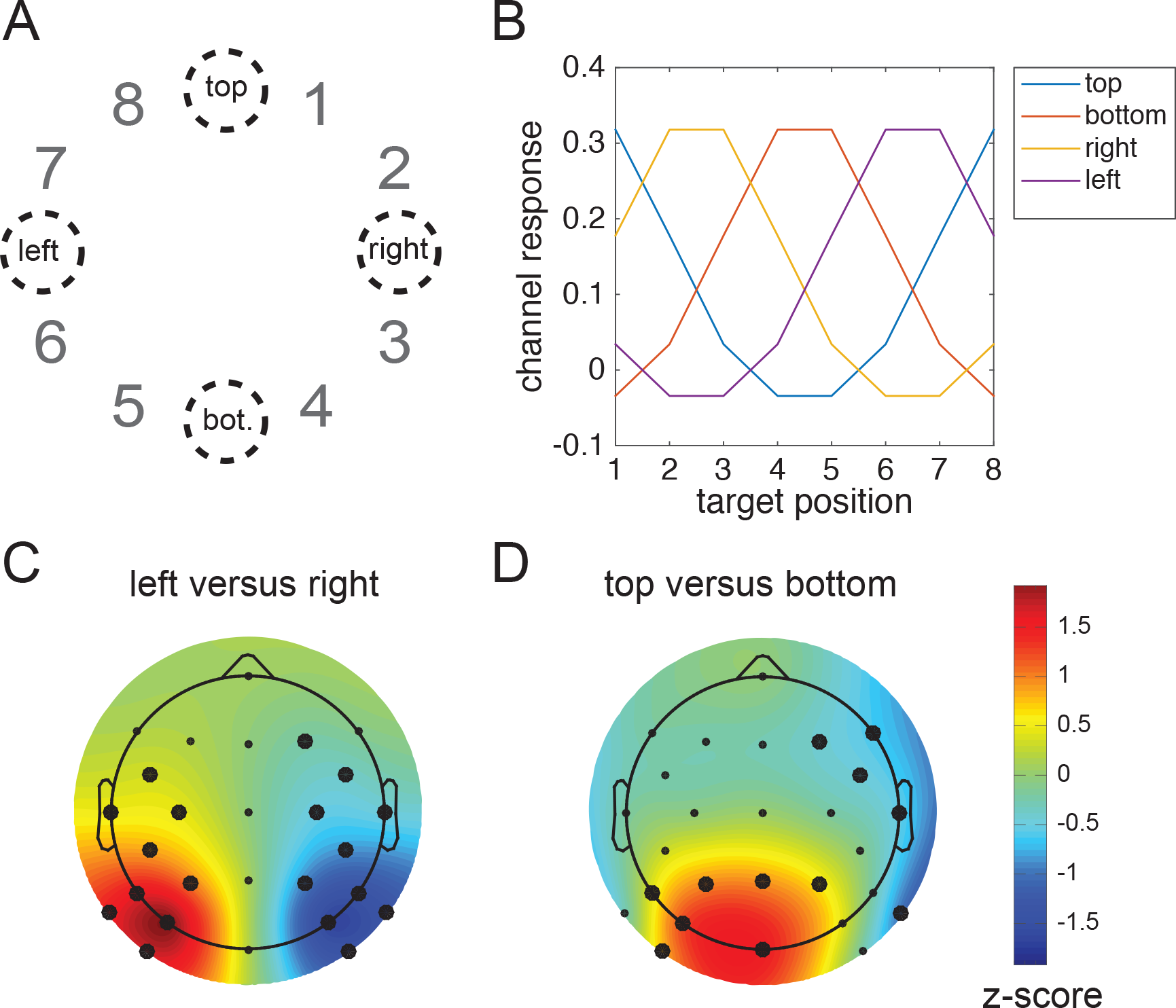
Reconstructing the neural signature of attentional capture for target positions that were never presented during Experiment 2. ***(A)*** The target positions that are reconstructed: top, bottom, left and right. ***(B)*** The constructed channel responses that are associated with these positions using the CTF from Figure 8A (see methods for details). Top is in between target position 8 and 1, so the channel response to top is constructed by averaging the hypothetical channel response to position 8 and position 1. Similarly, right is created from averaging channel responses to 2 and 3 etc. Any position on the circle can be constructed using a weighted average of channel responses. Left, right, bottom and top weights were reconstructed using the product of the constructed channel responses and the weight matrix at 260-270 ms. Next, the left versus right pattern was generated by subtracting the left from the right pattern ***(C)***, likewise the top versus bottom pattern was created by subtracting the bottom pattern from the top pattern ***(D)***. Note the similarity with the left-right and top-bottom patterns from Experiment 1. All patterns were normalized across space to allow direct comparisons. Thick electrode dots belong to a cluster that has p<0.01 under cluster based permutation testing.

Next, we multiplied these channel responses with the weights obtained from the initial training phase to construct four multivariate patterns that reflect the predicted hypothetical neural responses to targets at the top, bottom, left and right positions. To visualize the reconstructed response patterns to targets at the right versus left and targets at the top versus bottom locations, we subtracted the weights associated with the constructed right position from the weights for the constructed left position, and the weights for the constructed bottom position from the weights associated with the top position. The resulting values were normalized across space by subtracting the average over all electrodes and dividing by the standard deviation across electrodes. The topographic weight maps for hypothetical target locations on the horizontal midline (left versus right, both cluster-based p-values < 10^-3^ at 1000 iterations) and on the vertical midline (top versus bottom, cluster-based p-value < 10^-3^ for the positive cluster and cluster-based p-value = .009 for the negative cluster) are shown in Figures 9C and 9D. These two maps show remarkably similar topographies to the corresponding BDM-based maps for left versus right and top versus bottom targets obtained in Experiment 1 (Figure 5), where targets were in fact physically presented at these positions.

To formally evaluate this apparent match between constructed and real topographies associated with attended targets at left versus right or top versus bottom positions, we removed the four electrodes that were used in Experiment 2 but not in Experiment 1 from the data set for Experiment 2. We then correlated the group-averaged BDM-based activation patterns in Experiment 1 with the constructed activation patterns based on FEM weights in Experiment 2. In addition, we applied unpaired two-sided t-tests to compare BDM activation patterns and the FEM weights. These t-tests were performed for each electrode, correcting for multiple comparisons using cluster-based permutation testing. Both the left-right (Figure 10A) and the top-bottom topographies (Figure 10B) showed high correlations, and did not differ significantly when subtracted from each other. This strongly suggests that the reconstructed activity patterns obtained in Experiment 2 for hypothetical targets on the horizontal or vertical meridian showed the same topography as was found in Experiment 1 for real targets at these locations. To check whether the absence of significant differences between FEM weights and BDM patterns might be due to a lack of power (e.g. because we tested between rather than within participants), we also tested the left-right FEM weights against the top-bottom BDM pattern, this time yielding clear clusters of significant differences (p<10^-3^ for both clusters, two-sided cluster-based test at 1000 iterations, Figure 10C). This further confirms that the similarities between the reconstructed topographies found in Experiment 2 and the observed topographies found in Experiment 1 in Figure 10A and 10B are meaningful.

**Figure 10.**
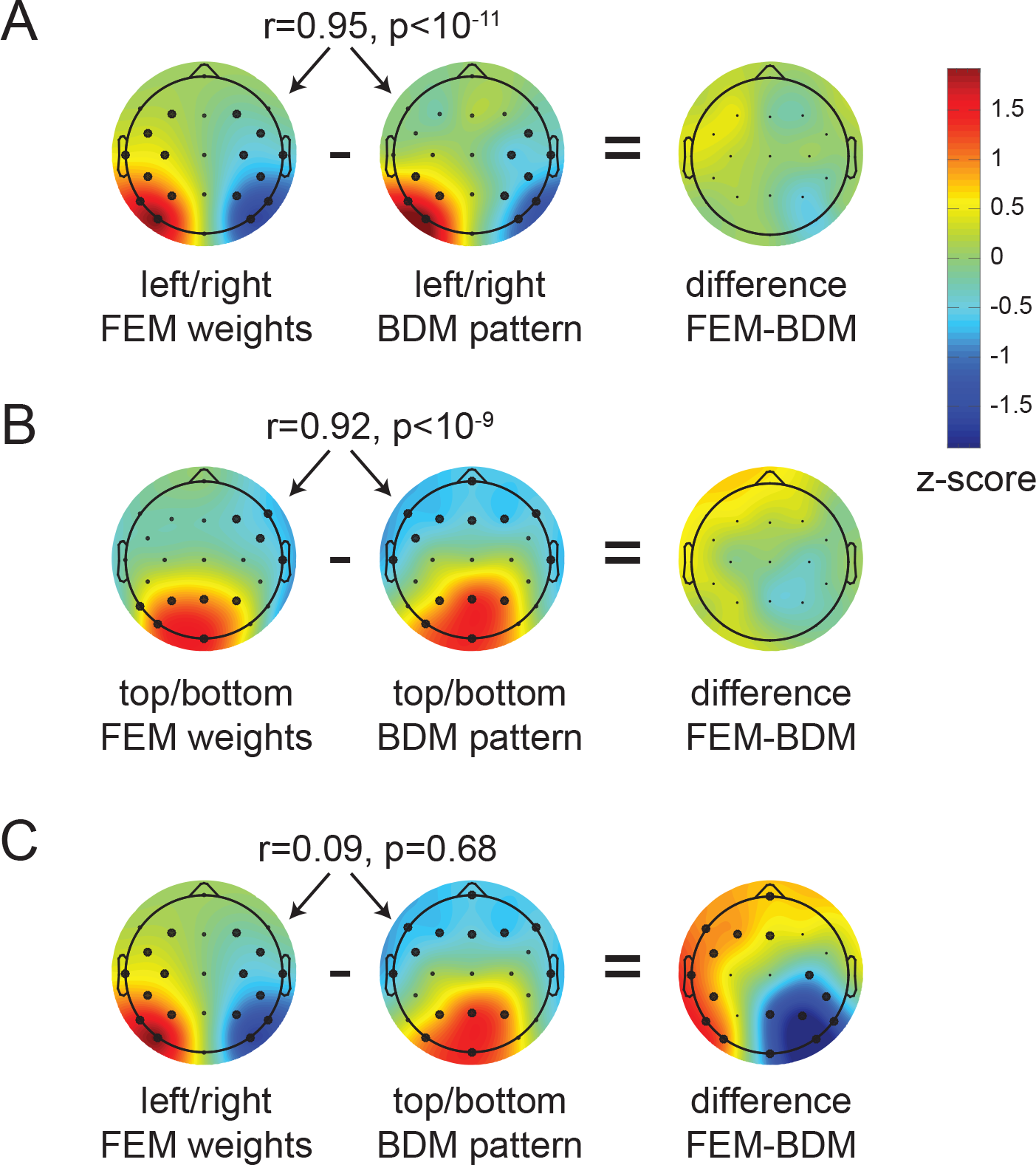
Comparing reconstructed weights in Experiment 2 to actual patterns in Experiment 1. The pearson correlation across population-averaged electrodes between FEM (Forward Encoding Model) weights from experiment 2 and BDM (Backward Decoding Model) patterns from experiment 1, as well as the unpaired t-test difference between FEM and BDM. ***(A)*** FEM and BDM left-right patterns show a high correlation and no significant differences. ***(B)*** FEM and BDM top-bottom patterns show a high correlation and no significant differences. ***(C)*** To show that the absence of significant differences in ***(A)*** and ***(B)*** is not caused by a lack of power due to t-testing between rather than within subjects, we also compare left-right FEM weights to the top-bottom BDM weights. These do not correlate, while their difference shows two large significant clusters. Thick electrode dots belong to clusters that have p<0.01 under cluster based permutation testing.

## Discussion

During search for targets with known features at unpredictable locations, feature-based selection mechanisms guide attention towards candidate target objects. To track these mechanisms, neural markers are needed that are not only temporally precise but can also dissociate selection at different locations within the visual field. Such markers can be provided by intracranial recordings, but these can only target few specific locations in the visual field a time, and are highly invasive. Many EEG and MEG studies^e.g. 5,7,25,26^ have used the N2pc component as a measure of feature-based selection^see^ ^8^ ^for review^. While the N2pc provides excellent temporal information, the fact that it is based on differences between ipsilateral and contralateral hemispheres severely constrains its ability to determine the spatial focus of feature-guided attention to discriminate between attended object locations in the visual field.

We demonstrate that a multivariate analysis of raw EEG data can track feature-based attentional selection in a spatially precise way while retaining the millisecond resolution inherent in EEG signals, exhibiting spatial selectivity from 200 ms onwards. In Experiment 1, backward decoding models successfully discriminated the selection of target objects on the vertical meridian (above versus below fixation) that is invisible to N2pc studies that measure attentional selection by comparing contralateral and ipsilateral ERP waveforms evoked by target objects in the left and right visual field. Classification accuracy rapidly increased from about 200 ms after display onset, thus matching the time course of the N2pc, and was highly comparable for top versus bottom and left versus right targets. In Experiment 2, classifiers successfully discriminated different target locations across all locations – even within the same quadrant – with classification performance again rising above chance level from about 200 ms post-stimulus. These results show that backward decoding models based on raw EEG data can be employed to obtain a much more precise spatial profile of feature-based attentional selection processes than is possible with standard N2pc methods.

We then employed a forward encoding model to construct a CTF that describes the relationship between target position and multivariate EEG in Experiment 2, analogous to recent studies of representations in working memory^14^. While in fitting the CTFs we used a delta basis set that does not make any a priori assumptions about their shape, the outcome revealed a clear gradual (i.e., Gaussian) response profile. This provides strong evidence that the spatial location of feature-based targets modulates the distribution of EEG activity in a continuous fashion.

Recent fMRI studies have used multivariate decoding to recover activation patterns in visual cortex that are sensitive to visual features or categories^e.g. 27-30^. In addition, forward encoding models have been developed for fMRI data^12,31^. Analogous multivariate analyses techniques have also been successfully applied to EEG data to track feature-selective activity in visual areas^13^ and the activation of stored items in visual working memory^32^. Two recent studies have demonstrated the utility of multivariate analyses of specifically EEG alpha band activity in tracking spatially selective processing during working memory maintenance and during cued shifts of spatial attention. Foster and colleagues asked participants to memorize the location of sample stimuli during delay periods, and found that the spatial distribution of alpha power tracked the sample stimulus locations in working memory^14^. Location-based CTFs based on multivariate alpha power showed a Gaussian profile, suggesting that location representations in working memory have a continuous relationship with the distribution of neural activity in visual cortex. Samaha and colleagues employed a spatial cueing task where covert attention had to be directed to one of six possible target locations indicated by centrally presented, symbolic cues^15^. Cued locations could be successfully classified on the basis of alpha activity elicited during the late period of the cue-target interval (1000 – 1900 ms after cue onset). Using a forward encoding model, Samaha and colleagues reconstructed CTFs, which revealed location-specific tuning that emerged around 450 ms after cue onset^15^. Here we employed analogous analyses to investigate feature-based attentional target selection in visual search, rather than endogenously cued spatial locations. Reliable tuning for target position emerged in the 200-300 ms time window in Experiment 2, which is much earlier than the onset of location-selective tuning reported by Samaha and colleagues (450 ms after cue onset)^15^. This difference suggests that the feature-based target selection mechanisms in this study are triggered more rapidly than endogenous spatial orienting processes elicited in response to the spatial cues, as in their study. Alternatively, our analyses are based on the raw EEG signal rather than time-frequency decomposed data, and may provide a more sensitive measure for such early signals. Moreover, the raw EEG signal has the additional benefit of retaining maximal temporal resolution, as it avoids the temporal blurring that inherently accompanies time-frequency analyses.

We believe one of the most exciting aspects of our data in combination with the forward modelling approach is that it allows for generalization beyond the observed. We used the graded CTF for the eight lateral target locations in Experiment 2 to reconstruct hypothetical channel responses and topographic activation patterns for target locations on the horizontal and vertical midlines that were not actually shown. We then compared these hypothetical patterns to the topographies of the BDM-derived neural activation patterns obtained in Experiment 1 on the basis of actual EEG responses to horizontal and vertical targets. The reconstructed topographies for these target locations were statistically indistinguishable from the neural activation profiles that were responsible for their successful decoding when they were physically present. The similarity between hypothetical and actual activation patterns is remarkable, given that separate sets of participants were tested. This shows that the FEM is even able to generalize to a different subject population. It further demonstrates the invertible nature of our FEM, and thus the continuous relationship between feature-based selection at particular locations and EEG activity patterns across electrodes. This shows promise for using these models as cortical mappers of feature-based attentional selection, analogous to the use of retinotopic maps of visual cortex obtained with fMRI measures^e.g. see^ ^33^. FEM-based methods may therefore have an important role for future studies of spatial attention and other types of cognitive operations in which a continuous relationship between cortical activity and some experimental variable exists.

We note that in the present experiments, above-chance classification emerged around 200 ms after search display onset but not earlier, which indicates that there were no EEG signals associated with target selection processes *prior* to the point in time when the N2pc typically emerges. This observation suggests that that the N2pc has probably rightly been regarded as the earliest electrophysiological marker of such processes. It is also notable that in both experiments, decoding accuracy remained above chance *after* the typical N2pc time window (beyond 300 post-stimulus). Similarly, the CTF constructed on the basis of our forward model in Experiment 2 showed reliable location tuning not only during the N2pc period, but also at longer latencies between 400 and 600 ms post-stimulus (see Figure 8C). These observations mirror previous ERP studies of target selection in visual search, which found a sustained contralateral posterior negativity (SPCN) during this time range^16,23,24^. The SPCN is usually interpreted as reflecting the encoding and processing of selected stimuli in visual working memory^e.g. 8,9^. Like the N2pc, SPCN components are based on activity differences between hemispheres, and therefore cannot reflect more subtle spatial patterns of neural activity associated with working memory maintenance within hemispheres. The current results suggest that multivariate decoding and encoding analyses can extract systematic patterns of EEG activity in the SPCN time range, and thus may enable more precise insights into how visual stimuli are spatially represented in working memory^14^, see also ^32^.

The decoding of attended target locations based on multivariate EEG data demonstrated in the current study has the potential to facilitate more general insights into the nature and time course of attentional selection. This method should be able to track shifts of attention from one candidate target object to another with high temporal precision, which could for example provide new information about the speed of such attention shifts. In displays where multiple candidate target objects are present, their respective locations may be decoded independently and simultaneously, in order to test whether and under which conditions attentional selection processes can operate in parallel at different locations in the visual field.

In summary, the current experiments demonstrate the potential of multivariate BDMs and FEMs of raw EEG data to overcome the limitations of the standard methodology that relies entirely on differences between cortical hemispheres. These new approaches to EEG analysis provide both spatially and temporally precise information about the spatial focus of feature-based attentional selection processes in visual search, and thus provide exciting new research opportunities.

## Acknowledgements

This work was supported by Open Research Area grant ES/L016400/1 from the Economic and Social Research Council (ESRC), UK, Open Research Area grant NWO 464-13-003, NL, and European Research Council Consolidator grant ERC-CoG-2013-615423.

## Author contributions

J.J.F., A.G., C.N.L.O. and M.E. designed the experiment, A.G. collected data,

J.J.F. analysed data, J.J.F., A.G., C.N.L.O. and M.E. wrote the paper.

